# Structures of the T cell potassium channel Kv1.3 with immunoglobulin modulators

**DOI:** 10.1101/2022.04.19.488765

**Authors:** Purushotham Selvakumar, Ana I. Fernández-Mariño, Nandish Khanra, Changhao He, Alice J. Paquette, Bing Wang, Ruiqi Huang, Vaughn V. Smider, William J. Rice, Kenton J. Swartz, Joel R. Meyerson

## Abstract

The Kv1.3 potassium channel is expressed abundantly on activated T cells and mediates the cellular immune response. This role has made the channel a target for therapeutic immunomodulation to block its activity and suppress T cell activation. We determined structures of human Kv1.3 alone, with a nanobody inhibitor, and with an antibody-toxin fusion blocker. Rather than block the channel directly, four copies of the nanobody bind the tetramer’s voltage sensing domains and the pore domain to induce an inactive pore conformation. In contrast, the antibody-toxin fusion docks its toxin domain at the extracellular mouth of the channel to insert a critical lysine into the pore. The lysine induces an active conformation of the pore yet blocks ion permeation. This study visualizes Kv1.3 pore dynamics, defines two distinct mechanisms to suppress Kv1.3 channel activity with exogenous inhibitors, and provides a framework to aid development of emerging T cell immunotherapies.

## INTRODUCTION

The Kv1.3 channel is a voltage-activated potassium (Kv) channel that is abundantly expressed on T cells where it plays an essential role in cell activation^1–4^. If Kv1.3 is inhibited or unable to conduct K^+^ ions, the T cell cannot undergo activation which in turn promotes immune suppression^5–12^. The ability to suppress the immune system via Kv1.3 motivates the development of therapeutic agents to inhibit the channel and ameliorate a number of autoimmune diseases including multiple sclerosis, type 1 diabetes, and rheumatoid arthritis^13^. The FDA-approved drug clofazimine inhibits Kv1.3 and is used for the treatment of psoriasis and graft-versus-host disease^14,15^, and other small molecule, peptide, and immunoglobin-based modulators are under development^13,16^. However, there are presently no reported structures of Kv1.3 in complex with any channel modulators, which limits our ability to rationalize their mechanisms.

Kv1.3 is one of eight members of the mammalian Kv1 family of Kv channels and the only one expressed on human T cells^1,2,17,18^. The Kv1 channels are tetramers and each subunit has a large cytoplasmic domain called T1, and six transmembrane helices called S1-S6, with S1-S4 forming a voltage-sensing domain (VSD) and S5-S6 forming the ion conducting pore domain^19–23^. The S5-S6 region harbors the P-loop, which in turn contains the highly conserved selectivity filter responsible for coordinating K^+^ ions and maintaining K^+^ permeation^24^. When Kv1 channels experience sustained activation they undergo C-type inactivation, a process that diminishes ion permeation and is thought to involve a conformational change in the selectivity filter that disrupts ion coordination^23,25–36^. Understanding the Kv1.3 channel gating process and how it can be pharmacologically manipulated is of fundamental importance to basic physiology and drug design.

Here we aimed to define the mechanisms for two recently developed immunoglobulin-based Kv1.3 modulators, a nanobody and an engineered IgG antibody. First, we solved the cryo-electron microscopy (cryo-EM) structure of human Kv1.3 without modulators as a reference. We find the selectivity filter adopts two distinct conformations which are both consistent with an inactive channel state expected from the experimental conditions, and different from previously reported Kv1 family structures which were in active state conformations^20,23,36,37^. Next, we solved the structure of Kv1.3 with the recently-developed nanobody inhibitor A0194009G09 (Ablynx/Sanofi) which blocks with nanomolar affinity through an unknown mechanism and has 1,000-fold selectivity for Kv1.3 over hERG, Kv1.5, and Kv1.6^38^. We then solved the structure of Kv1.3 with MNT-002 (Minotaur Therapeutics), an engineered antibody where the peptide toxin ShK was fused into the complementary determining region 3 (CDR3) of an IgG antibody. ShK or its derivatives block Kv1.3 with high selectivity, greater than 100-fold over other Kv channels^39,40^, and show efficacy in rat models of autoimmune conditions including arthritis^10,41^, diabetes^42,43^, dermatitis^44^, and asthma^45^. The Fc region of the antibody engages Fc receptors to enhance inhibition of T cells (manuscript in preparation). Taken together these results enrich our understanding of Kv1.3 pore dynamics and channel gating, and establish mechanisms of Kv1.3 modulation by two distinct molecules.

## RESULTS

### Structure of human Kv1.3 and its selectivity filter conformations

Before investigating Kv1.3 in complex with modulators we first aimed to define a high-resolution reference structure of the channel in an unbound state. We expressed and purified full-length human Kv1.3 and solved the structure using single particle cryo-EM to 2.89 Å with C4 symmetry (**Fig. 1A, Fig. S1, Table S1**). Inspection of the density map showed clear densities for the VSDs, the channel pore domain, the cytoplasmic T1 domains, and the caveolin-binding domains^46^ (**Fig. 1A, 1B, Fig. S1**). The overall architecture of the structure is consistent with a recent structure of Kv1.3^47^, structures of Kv1.2^19,20^, the Kv1.2-2.1 chimera^21,22,36^, and the orthologous Shaker Kv channel from *D. melanogaster*^23^. Putative detergent densities were observed at the hydrophobic interior surface of each S6 helix but show no apparent influence on the structure (**Fig. S1**). The VSDs are in the ‘up’ position (**Fig. 1C**) and the internal gate is open (**Fig. 1D, Fig. S4**), reflecting the fact that in the experimental conditions the channel is at 0 mV and not at a negative voltage which would otherwise keep the channel in a resting conformation with the VSDs ‘down’ and the internal gate closed.

**Figure 1.**
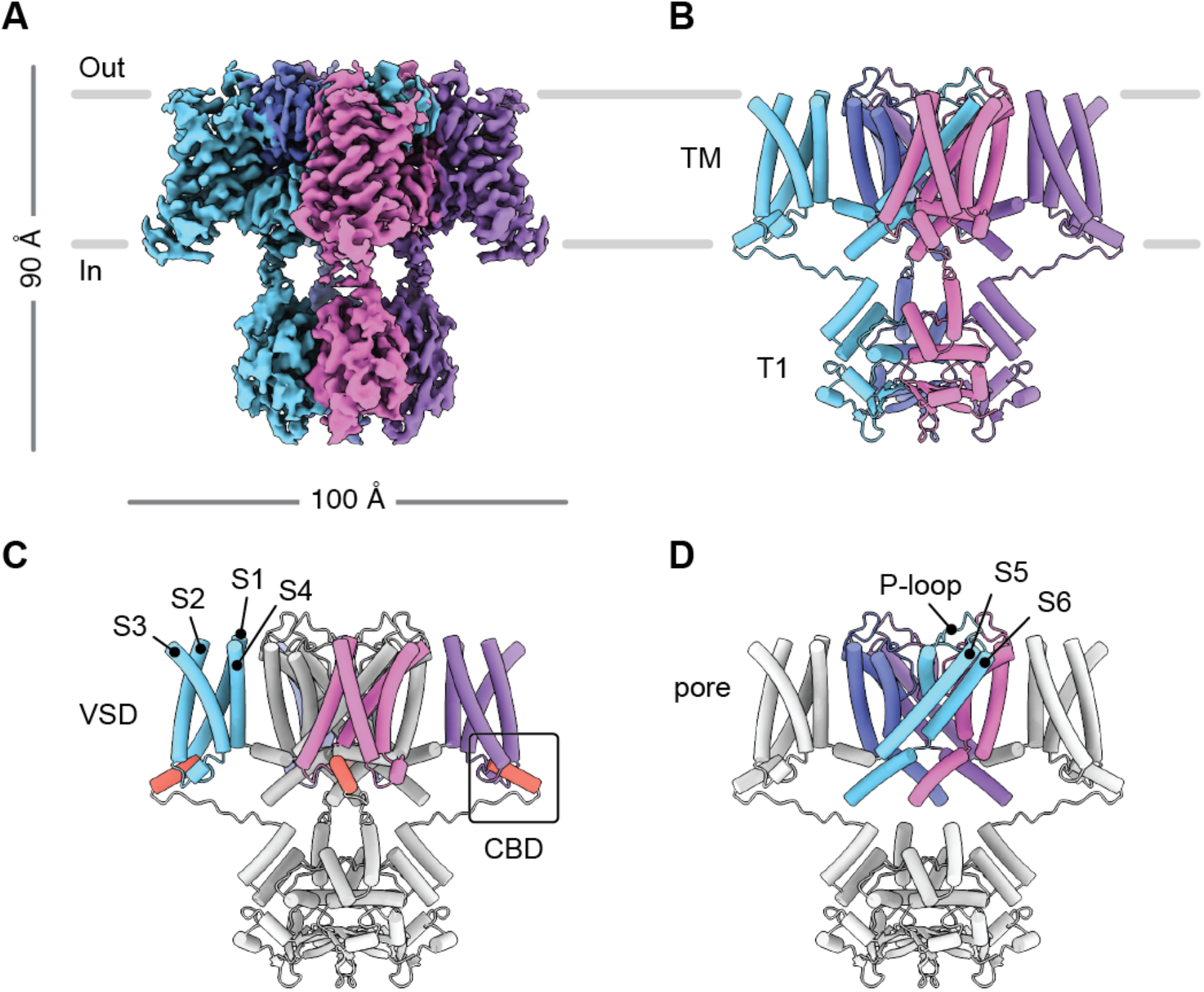
Structure of human Kv1.3. **(A, B)** Cryo-EM map (A) and model (B) of human Kv1.3. The transmembrane (TM) region and T1 domains are indicated. **(C, D)** Model of Kv1.3 with the voltage sensing domains (VSDs) and caveolin binding domains (CBDs) colored as indicated (C), or with the pore domain colored as indicated (D). The front and rear VSDs are omitted in for clarity in (D).

Next, we inspected the pore region of the cryo-EM map and evaluated the conformation of the selectivity filter (**Fig. 2**). We observed the P-loop appears to adopt two discernable conformations in the short region from Tyr447 to Met450 (**Fig. 2A, 2B**). The conformations were present in all four subunits even when the data was processed without imposing symmetry (C1), suggesting that our initial particle classification and structural refinement was not sensitive to this particular structural variation. These observations motivated us to obtain separate structures for each loop conformation using computational image classification, and also address whether individual tetramers are conformationally homogeneous in the loop region or if tetramers contain a combination of the two subunit conformations. We used symmetry expansion in Relion^48^ to isolate each subunit within each tetramer and 3D classification to obtain density maps for the Kv1.3 subunit in both conformations (**Fig. 2C, Fig. S1**). The Kv1.3 structure was interpreted and modeled in two conformations using the C4 symmetric tetramer map in tandem with the individual subunit maps.

**Figure 2.**
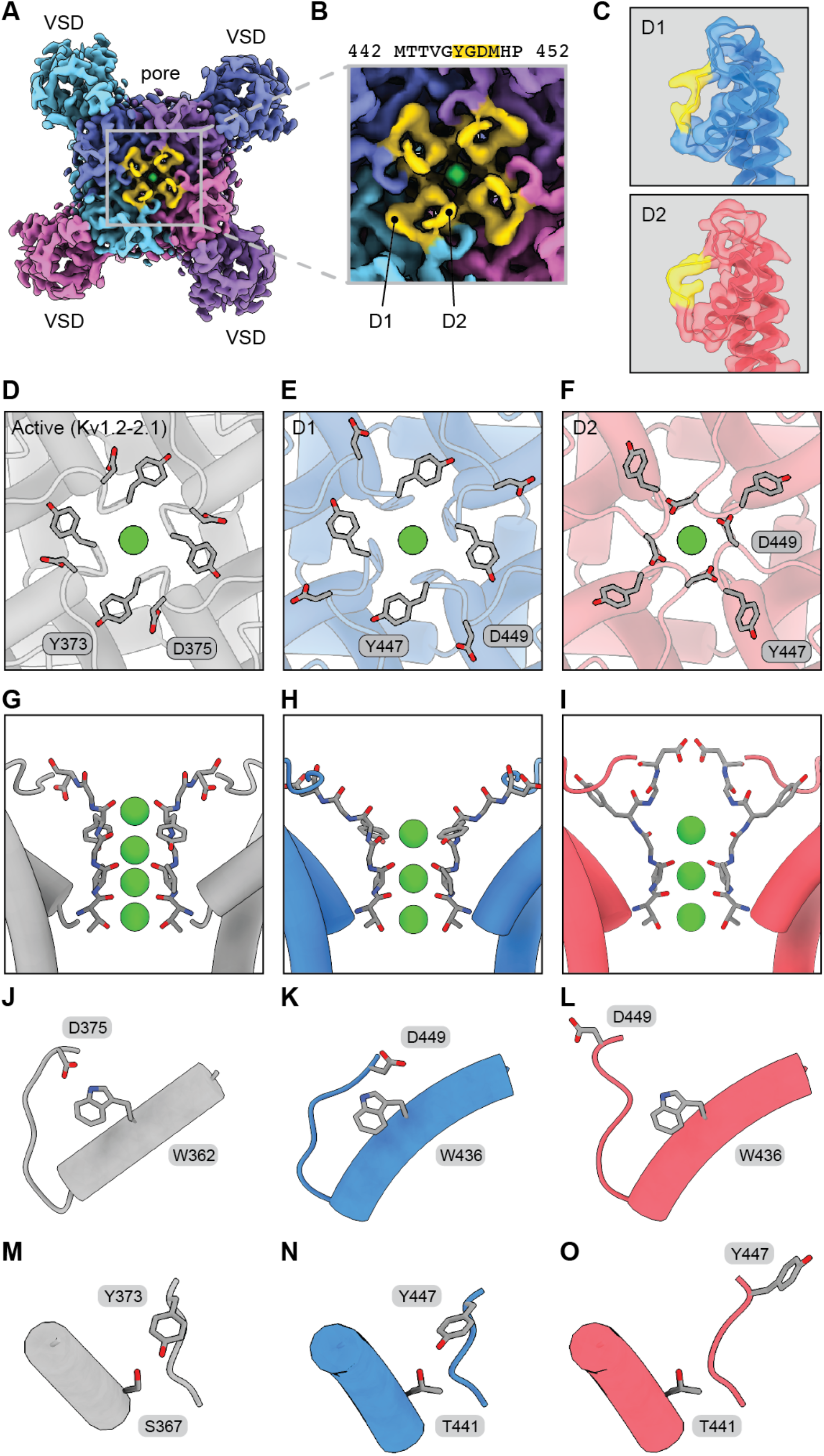
The Kv1.3 selectivity filter adopts two distinct dilated conformations. **(A, B)** Kv1.3 cryo-EM map as viewed from the extracellular space with the D1 and D2 loop regions highlighted in yellow. Panel (A) shows the entire extracellular surface of the channel and (B) shows a magnified view of the pore region. **(C)** Cryo-EM maps for individual D1 and D2 subunits as determined by symmetry expansion and 3D classification. The maps are shown with models to highlight the different selectivity filter conformations. The region from Tyr447 to Met455 is highlighted in yellow. **(D-I)** Views of the pore for active state (Kv1.2-2.1, PDB 2R9R), D1 conformation, and D2 conformation. The pores are shown from the extracellular space (D-F) or from the side perspective (G-I). **(J-O)** Asp and Trp residues (J-L) and Tyr and Ser/Thr residues (M-O) involved in hydrogen bonds that regulate C-type inactivation.

To analyze the Kv1.3 subunit conformations we required a reference structure in an active state and selected the Kv1.2-2.1 structure^21^ (**Fig. 2D**). This structure was chosen because of its sequence similarity to Kv1.3, and its high resolution compared to a recent Kv1.3 structure which was of insufficient resolution to confidently interpret its pore conformation and ion occupancy^47^. Both of our Kv1.3 subunit conformations show a dilation of the selectivity filter with K^+^ ion densities in three ion binding sites (**Fig. 2E, 2F, Fig. S1, S4**). In contrast, the active state reference structure has a narrow selectivity filter coordinating four K^+^ ions (**Fig. 2D**). Accordingly we designated the two subunit conformations in our Kv1.3 structure dilated 1 (D1) and dilated 2 (D2) (**Fig. 2E, 2F**). The first dilated conformation (D1) shows the apex of the P-loop folded outward relative to the reference structure (**Fig. 2G, 2H**). In this conformation, Asp449 of Kv1.3 is directed upward and away from the channel, and in particular is displaced from Trp436 (**Fig. 2K**). This is significant because in the active state structure the corresponding Asp (Asp375) orients downward with its carboxylate moiety positioned near the indole nitrogen of Trp (Trp362) (**Fig. 2J**). The second dilated conformation (D2) shows a more severe disruption of the selectivity filter (**Fig. 2I**). Asp449 is displaced from Trp436 in D2, just as in D1, but Asp449 adopts an entirely different position in D2 (**Fig. 2L**). In addition, Tyr447 points away from Thr441 (**Fig. 2O**) which is important because these two residues are hydrogen bonding partners in the active reference structure (**Fig. 2M**) and the D1 conformation (**Fig. 2N**). The Asp-Trp and Tyr-Thr hydrogen bonds are highly conserved structural motifs in Kv channels, and rupture of the bonds is thought to accompany C-type inactivation in the Shaker Kv channel^25,27,33,49,50^. Also significant is that the D2 conformation of the Kv1.3 selectivity filter resembles structures of mutants of the *D. melanogaster* Shaker Kv channel^23^ and the Kv1.2-2.1 chimera ^36^ which are proposed to be C-type inactivated conformations.

The 3D subunit classification analysis provided conformational identities of individual subunits in the dataset, which in turn enabled a mapping of these identities back to the parent tetramers (**Fig. S1, Table S2**). In this way we determined the conformational composition of each tetramer. We excluded from the analysis those tetramers for which one, two, or three subunits could not be classified. This left 6,598 tetramers where all four subunits could be identified as either D1 or D2 (**Table S2**). Out of these tetramers, the majority were a mixture of D1 and D2 subunits with 1:3 (D1:D2) being the most abundant stoichiometry (30.8% of tetramers), followed by 2:2 (24.5%) and 3:1 (13.6%) (**Table S2**). The remaining tetramers were either pure D1 having a 4:0 ratio (4.5%) or pure D2 with a 0:4 ratio (26.6%) (**Table S2**). Together this analysis showed that the vast majority of tetramers are a mixture of D1 and D2 subunits, suggesting that the D1 and D2 filter conformational changes do not occur in a fully cooperative manner.

### The nanobody binds and bridges the voltage sensing and pore domains

We first validated the function of purified A0194009G09 nanobody (NB) by applying it to cells expressing Kv1.3 and measuring K^+^ currents (**Fig. 3A-3C**). These current recordings showed that under control conditions, Kv1.3 was activated by membrane depolarization and displayed its characteristic slow C-type inactivation. In contrast, when NB (100 nM) was added to the extracellular solution Kv1.3 channel inactivation was greatly accelerated and channel current was almost completely inhibited by the end of the test depolarizations (**Fig. 3A-3C**). The structure of Kv1.3 with the NB inhibitor was resolved at 3.25 Å resolution and four copies of the NB were observed on the extracellular surface of Kv1.3 (**Fig. 3D-3F, Fig. S2**). Inspection of the structure showed that only three K^+^ ions are present in the pore (**Fig. 3G, Fig. S4**), as seen in the D1 and D2 conformations from the unbound structure (**Fig. 2**). Similar to the D1 conformation, the NB-bound structure shows Asp449 flipped away from Trp436 in an orientation incompatible with hydrogen bond formation (**Fig. 3H**). However, Tyr447 adopts two different rotamers in the density map. One Tyr447 rotamer matches D1 with its hydroxyl group hydrogen-bonded to Thr441 (**Fig. 2N, 3H**). The second rotamer has its hydroxyl group directed towards the extracellular space and so it is distinct from both D1 and D2 (**Fig. 3H**). For this reason, we designated the conformation with the second Tyr447 rotamer as D3. Thus, when Kv1.3 is bound by the nanobody the selectivity filter adopts D1 and D3 conformations.

**Figure 3.**
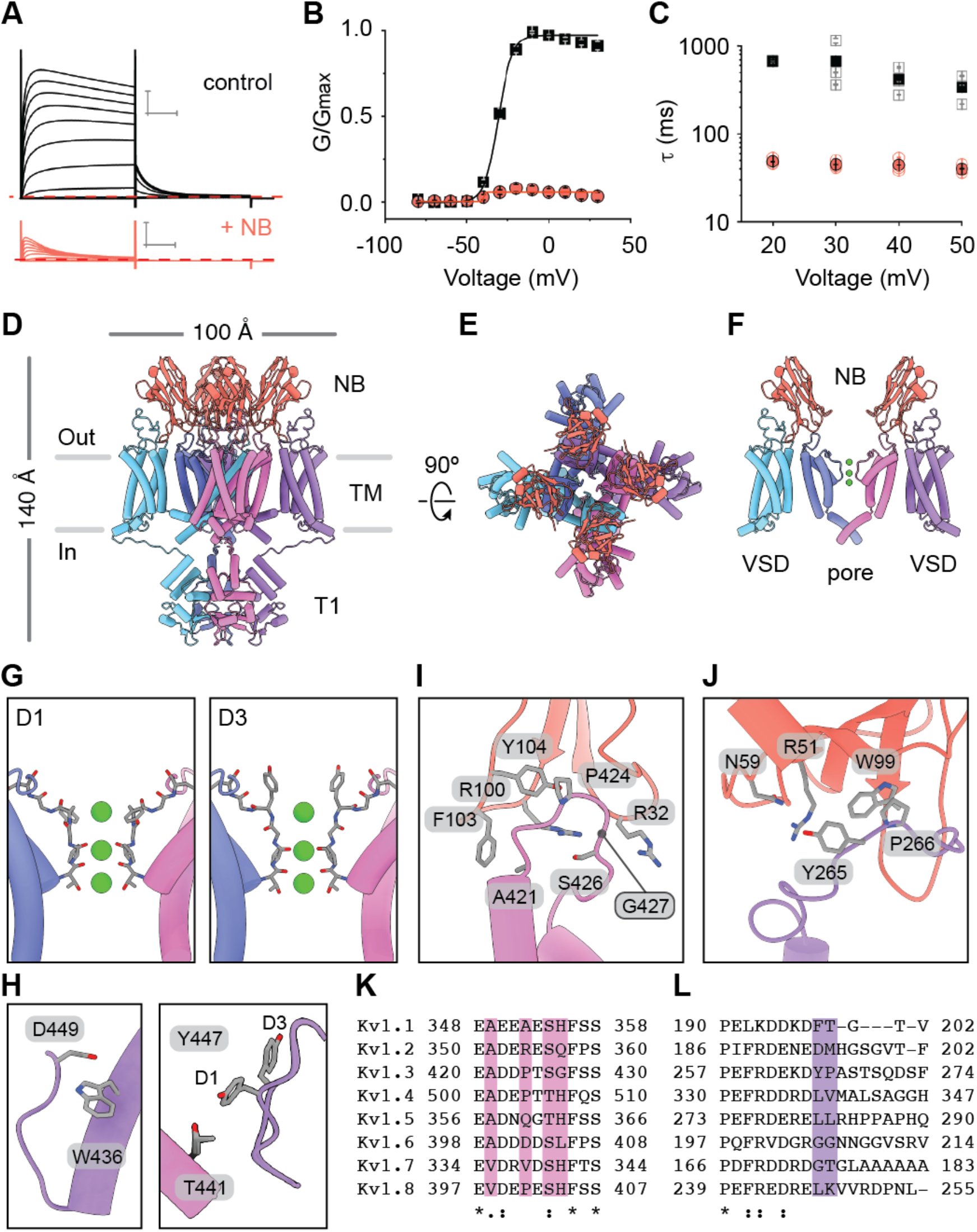
Multiple nanobodies bind the turret loops and voltage sensing domains of Kv1.3. **(A)** Kv1.3 current traces obtained from a family of depolarizing pulses ranging from −80 mV to +30 mV in control conditions (black) and in the presence of 100 nM nanobody (NB) A0194009G09 (orange). Holding voltage was −80 mV and tail voltage was −50 mV. Red dotted line denotes zero current level. Scale bar indicates 2 μA and 50 ms. **(B)** G-V relations obtained in control solution (black squares; n=7) or 100 nM nanobody (orange circles; n=7) by measuring the peak of the tail current at −50 mV and normalizing it to the maximum tail current in control solution. Solid lines are fits of the Boltzmann equation to the data and error bars are SEM. **(C)** Time constants of inactivation measured from individual experiments for test depolarizations to between +20 and +50 mV (control: open black squares, 100 nM nanobody: open orange circles) and averages (control: black filled squares, 100nM: orange filled circles) (n=3) where error bars are SEM. **(D, E)** Side (D) and extracellular (E) views of the model for Kv1.3 in complex with four nanobodies (red). **(F)** Cutaway view of the structure in (D) showing two nanobodies with each bound to a pore domain and a VSD of Kv1.3. The T1 domains are omitted for clarity. **(G)** Structures of the selectivity filter of the Kv1.3-nanobody complex with the D1 (left) and D3 (right) conformations. **(H)** Views of W436/D449 (left) and T441/Y447 (right) for both the D1 and D3 conformations identified in the Kv1.3-nanobody structure. The panel with T441/Y447 shows the D1 and D3 models superimposed to highlight the different rotamers for Y447. **(I, J)** Close up views of a nanobody in complex with a turret loop of a pore domain (I) and the S1-S2 loop of a VSD (J). **(K, L)** Sequence alignment for the turret loop (K) and VSD S1-S2 loop (L) regions for human Kv1 channels.

The nanobodies do not make direct contact with the selectivity filter of Kv1.3, but rather they decorate the turret loops near the pore and extend contacts to the VSDs (**Fig. 3F, 3I, 3J**). On the turret loop, Phe103 of the NB forms a hydrophobic interaction with Ala421, R100 forms a hydrogen bond with Ser426, and Tyr104 and Pro424 make hydrophobic contact (**Fig. 3I**). Arg32 of the NB extends directly over Gly427 of the turret loop. This glycine is notable because all other human Kv1 channels display large side chains (His, Gln, or Leu) at the equivalent positions, which may clash with Arg32 (**Fig. 3I, 3K**). Arg51 and Asn59 of the NB both appear capable of hydrogen bonding with Tyr265 on the VSD, while Trp99 of the NB forms a hydrophobic interaction with Pro266 (**Fig. 3J, 3L**).

The structure shows four NBs can bind the Kv1.3 tetramer (**Fig. 3D, 3E**) but it was solved with a threefold molar excess of NB (three NB per Kv1.3 subunit). This motivated us to investigate the dose response of Kv1.3 to NB. First, we measured the rate of inactivation without NB or with 10 or 100 nM NB. Inactivation with no NB could be described by a single exponential function and was relatively slow (**Fig. 3C, Fig. S5A, S5B**). With 10 nM nanobody inactivation was accelerated and was modeled with multiple exponential functions (**Fig. S5A, S5B**), while at 100 nM inactivation was over 10-fold faster (**Fig. 3C, Fig. S5A, S5B**). Second, we examined the concentration dependence for the onset and the recovery from inhibition by the NB. Onset of inhibition was sped by increasing the concentration of the NB from 10 nM to 100 nM (**Fig. S5**). In contrast, recovery from inhibition was slowed by increasing the NB concentration. This was inferred from the sigmoidal recovery profile following removal of the NB, and the increase in this sigmoidicity with higher NB concentration (**Fig. S5**). Taken together, these functional results show that NB binding occupancy on Kv1.3 can vary and that higher occupancies promote faster and more durable inactivation.

### ShK toxin block induces a conductive pore conformation

To investigate the MNT-002 antibody-ShK fusion we generated the Fab-ShK form of the molecule and validated its ability to block Kv1.3 currents (**Fig. 4A, 4B**). We then bound Fab-ShK to Kv1.3 and solved the structure to a global resolution of 3.39 Å. The density map readily enabled atomic modeling of Kv1.3 and the ShK toxin domain (**Fig. 4C, Fig. S3**). However, the Fab region was only weakly resolved so was modeled by rigid body fitting a homology model into the density map (**Fig. 4C, Fig. S3**).

**Figure 4.**
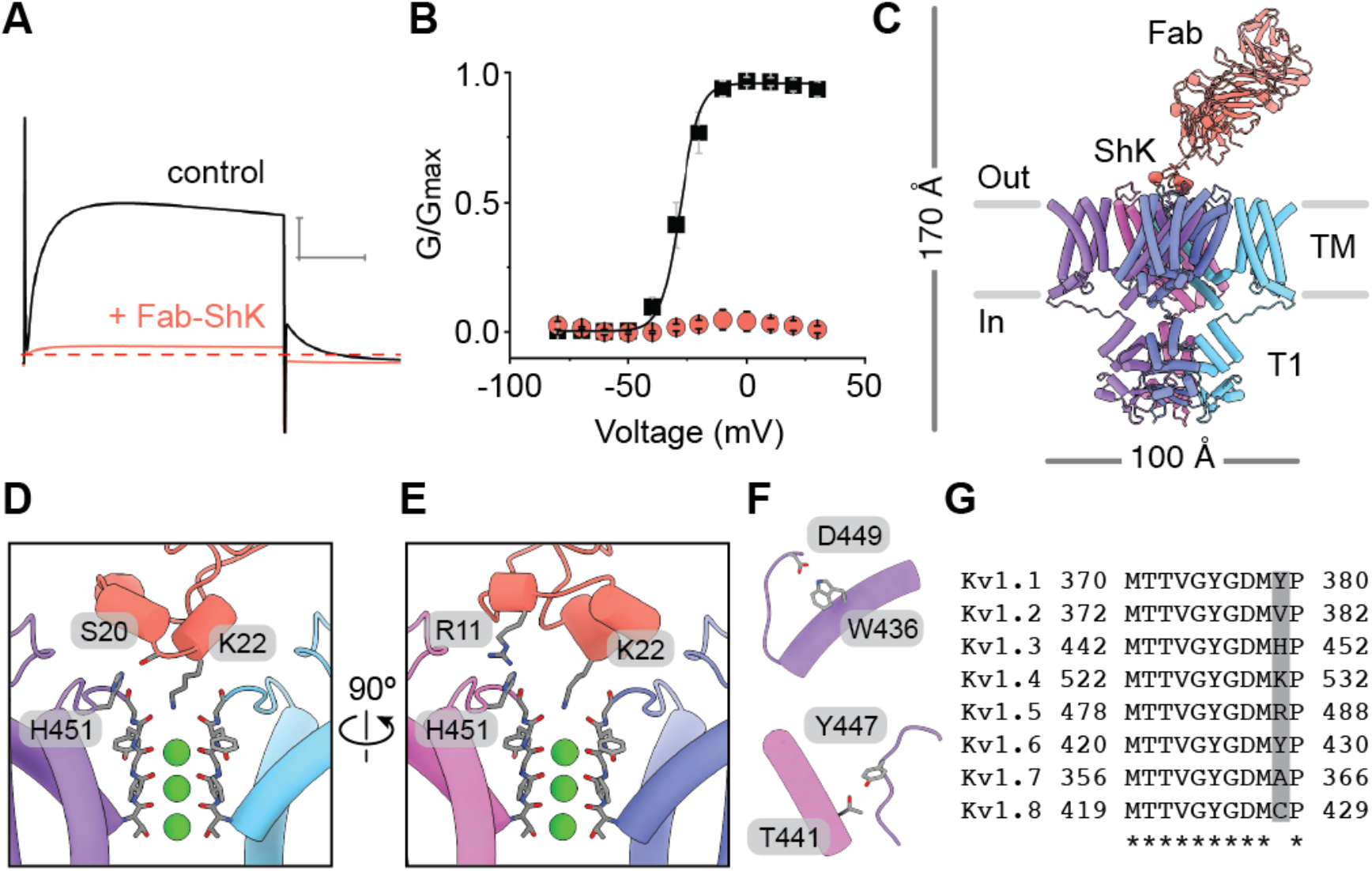
Fab-ShK induces a conductive pore conformation to block the channel. **(A)** Kv1.3 current traces obtained by depolarizations to 0 mV from control solution (black) and in the presence of 10 nM ShK (orange). Holding voltage was −80 mV and tail voltage was −50 mV. Red dotted lined denotes zero current level. Scale bar indicates 5 μA and 50 ms. **(B)** G-V relations obtained in control solution (black squares; n=5) or 10 nM ShK (orange circles; n=5) by measuring the peak of the tail current at −50 mV and normalizing it to the maximum tail current in control solution. Solid lines are fits of the Boltzmann equation to the data and error bars are SEM. **(C)** Model of Kv1.3 in complex with Fab-ShK as viewed from the side. **(D, E)** Close up views of ShK docking at the selectivity filter of Kv1.3. The ShK residue numbers correspond to ShK alone, not their numbering in the Fab-ShK fusion. The residues Arg11, Ser20, and Lys22 of ShK correspond to Arg118, Ser127, and Lys129 in the Fab-ShK fusion. **(F)** Key hydrogen bonds involved in channel gating are stabilized when Kv1.3 is bound by ShK. **(G)** Sequence alignment for the selectivity filters of Kv1 channels with the position for His451 of Kv1.3 highlighted.

The Fab binds with an ~45° angle to the extracellular surface of Kv1.3 and its long and narrow CDR3 ‘stalk’ allows it to avoid contact with the extracellular loops of Kv1.3 and position ShK at the mouth of the pore (**Fig. 4C**). ShK forms its most significant contact with the pore via Lys22, with the ammonium group hooked downward into the selectivity filter to coordinate the backbone carbonyl of Tyr447 on all four Kv1.3 subunits (position TVGYG of the K^+^ channel signature sequence) (**Fig. 4D, 4E, Fig. S4**). In this way Lys22 functions as a surrogate K^+^ ion as proposed^51–54^. We compared the conformation of the selectivity filter in the Kv1.3 with Fab-ShK structure to the active state reference structure and the D1 and D2 conformations observed in unbound Kv1.3 (**Fig. 2**). This showed that when the selectivity filter is bound by ShK it adopts a similar conformation as seen in active K^+^ channel structures^21,23,36^ (**Fig. 2J, 2M, 4D, 4E**). Despite the absence of the outermost K^+^ ion in the filter, the key hydrogen bonds formed by Asp449/Trp436 and Y447/T441 are intact (**Fig. 4F**), consistent with an active channel conformation (**Fig. 2J, 2M**). Thus, ShK induces an active conformation of the selectivity filter despite blocking ion flow. In addition to the contact between Lys22 and the selectivity filter, Ser20 and Arg11 of ShK form hydrogen bonds with His451 of Kv1.3, but on separate subunits (**Fig. 4D, 4E**). This is significant because Ser20 and Arg11 are major determinants of ShK toxin binding^52^ and because Kv1.3 is the only family member with a histidine in this position, with most other Shaker channels having residues that would be less suited to forming the contact with Ser20 because of charge or steric mismatch (**Fig. 4G**).

## DISCUSSION

In this study we determined the mechanisms of Kv1.3 modulation for a recently-developed nanobody inhibitor and an engineered fusion of an IgG antibody with the ShK toxin. We first solved a structure of unbound Kv1.3 which showed a dynamic selectivity filter with two discernable conformations (D1, D2) (**Fig. 1, 2)**. The dilation of the filter in D1 and D2 reorients the critical backbone carbonyls to disrupt K^+^ ion coordination, a feature which has been proposed to be associated with C-type inactivation^23,36^ and may serve to increase the energetic barrier to K^+^ permeation^55^. The dilated conformations also show disruption of two key hydrogen bonds that are thought to stabilize an active channel conformation (**Fig. 2)**^27,33,49,50,56^.

The A0194009G09 nanobody-bound structure reveals an unanticipated mechanism of channel inhibition in which the nanobody bridges the turret loops and the VSDs to promote C-type inactivation (**Fig. 3, Fig. S5)**. The structure has two distinct conformations (D1, D3) distinguished by different rotamers for Tyr447 which is a critical residue in C-type inactivation^33^. It is notable that the regions where the nanobody binds are among the most variable segments in the Kv1 channel family (**Fig. 3**). Atomic delineation of the nanobody interaction with Kv1.3, along with increasing availability of potassium channel structures, opens the possibility of designing new nanobodies with high specificity for different channels. The FDA approval in 2019 of the first nanobody drug^57^ and the exponential rise of FDA-approved immunoglobulins^58^ suggests this may be a productive direction for future work.

The structure of Kv1.3 with the Fab form of the MNT-002 antibody shows the ShK toxin switches the selectivity filter from the D1 and D2 conformations present in the unbound structure to an active conformation, yet blocks the ion permeation pathway (**Fig. 4)**. ShK is a prominent member of a large family of sea anemone toxins that bind many K^+^ channels^59^ so the structure serves as a template to understand these toxins. In addition, many animal venom toxins block K^+^ channels and have been extensively studied, but the structural details of these interactions have been challenging to fully define owing to methodological limitations^51^. The venom toxins, like ShK and the sea anemone blockers, employ a highly conserved lysine residue to block the channel. Thus, the details of our structure may inform the understanding of channel block for many classes of toxins and potassium channels.

## ACKNOWLEDGEMENTS

We thank Tsg-Hui Chang for frog keeping and surgery, Abigail Kelley for technical support, and members of our labs, Alessio Accardi’s lab and Olga Boudker’s lab for discussion. Cryo-EM was performed at the NYU Langone Health’s Cryo-Electron Microscopy Laboratory, the National Cancer Institute’s National Cryo-EM Facility (NCEF) at the Frederick National Laboratory for Cancer Research under contract HSSN261200800001E, and at the Simons Electron Microscopy Center and National Resource for Automated Molecular Microscopy located at the New York Structural Biology Center and supported by grants from the Simons Foundation (SF349247), NYSTAR, and the NIH National Institute of General Medical Sciences (GM103310). Assistance with cryo-EM was provided by Tara Fox and Thomas Edwards (NCEF), and Eugene Chua and Edward T. Eng (New York Structural Biology Center). Molecular graphics were prepared and analysis performed with UCSF ChimeraX, developed by the Resource for Biocomputing, Visualization, and Informatics at the University of California, San Francisco with support from National Institutes of Health R01-GM129325 and the Office of Cyber Infrastructure and Computational Biology, National Institute of Allergy and Infectious Diseases. This work was supported by NIH grant R01 GM105826 to V.V.S., National Institute of Neurological Disorders and Stroke NS002945 to K.J.S., and a Leon Levy Fellowship in Neuroscience grant and a STARR Cancer Consortium grant to J.R.M.

## AUTHOR CONTRIBUTIONS

PS purified Kv1.3 and the A0194009G09 nanobody; RH purified the MNT-002 Fab; PS, AP, BW, and WJR performed cryo-EM; PS, NK, CH, and JRM performed structure determination; AIFM performed electrophysiological experiments; PS and JRM wrote the manuscript; all authors contributed to manuscript editing and preparation.

## METHODS

### Expression and purification of human Kv1.3

The gene for human Kv1.3 was cloned into the pEZT-BM BacMam expression vector^60^ and at the C-terminus was fused to a 3C protease recognition site followed by the mVenus fluorescent protein and a Twin-Strep affinity tag. The expression construct was transformed into DH10Bac cells to produce bacmid which was then transfected into Sf9 cells grown in ESF 921 media (Expression Systems). P1 and P2 virus production was monitored using GFP fluorescence from the pEZT-BM vector until virus harvesting. HEK293S GnTI-cells (ATCC CRL-3022) were grown (3.2 L or 6.4 L) at 37°C and 8% CO2 to a density of 3.5×10^6^ cells/mL in FreeStyle suspension media (Gibco) supplemented with 2% fetal bovine serum (Gibco) and Anti-Anti (Gibco). P2 virus was added to cells, the suspension was incubated at 37°C for 24 hrs, then sodium butyrate (Sigma) was added to a final concentration of 10 mM and flasks were shifted to 30°C and 8% CO2. Cells were collected 84 hrs after transduction by low-speed centrifugation, flash-frozen in liquid nitrogen, and stored at −80°C.

Cell pellets were resuspended in ice-cold resuspension buffer containing 50 mM Tris (pH 7.5), 150 mM KCl, 2 mM DTT, protease inhibitor tablet (Sigma), 1 mM PMSF, 0.5 mM EDTA, and 25 μg/ml DNAse (6 mL buffer per 1 g of cell pellet) and manually pipetted until no clumps remained. The cells were disrupted on ice using a QSonica Q700 sonicator (3×15 sec, power level 60), lysate was clarified by low-speed centrifugation (7,200×g for 20 min), and membranes were isolated by ultracentrifugation at 125,000×g for 2 hrs. Membranes were resuspended and homogenized in buffer containing 20 mM Tris (pH 7.5), 150 mM KCl, protease inhibitor tablet (Sigma), 1 mM PMSF, and 0.5 mM EDTA. The protein was extracted by adding 50 mM n-Dodecyl-β-D-maltopyranoside with 0.25% cholesteryl hemisuccinate (Anatrace, D310-CH210) and nutating for 1 hr at 4°C. The mixture was ultracentrifuged at 125,000 × g for 50 min to remove insoluble material. The supernatant was filtered through a 0.45 μm filter then bound to a 5 mL Strep-Tactin column (GE) equilibrated with running buffer containing 20 mM Tris (pH 7.5), 150 mM KCl, 0.5 mM EDTA, and 2 mM DDM with 0.01% CHS. The bound protein was washed with running buffer containing 10 mM MgCl_2_ and 5 mM ATP to remove bound heat shock proteins ^61^, then eluted with running buffer composed of 10 mM desthiobiotin (IBA). Elution fractions were collected, analyzed by SDS-PAGE, concentrated, and loaded onto a Superose 6 Increase 10/300 GL (Cytiva) column equilibrated in buffer containing 20 mM Tris (pH 7.5), 150 mM KCl, 2 mM DTT, 0.5 mM EDTA, and 0.5 mM DDM with 0.0025% CHS. Elution fractions were collected, analyzed by SDS-PAGE, and peak fractions were concentrated and used for cryo-EM experiments.

### A0194009G09 nanobody expression and purification

The gene sequence for nanobody A0194009G09 was obtained from patent CA2951443A1^38^, codon optimized for *E. coli* expression, then synthesized and cloned into the pET26b(+) vector in frame with an C-terminal 6 × His tag (GenScript). BL21 DE3 cells were transformed with the plasmid and grown at 37°C in TB media supplemented with 1 mM MgCl_2_, 0.01% glucose, and kanamycin (25 μg/ml). When the cells reached OD600 of 0.7, 1 mM IPTG was added to induce protein expression. After 3 hrs the cells were harvested and pellets stored at −80°C. To purify the protein, cell pellets were thawed and added to buffer containing 20 mM Tris (pH 7.5), 150 mM KCl,1 mM PMSF, 5 mM MgCl_2_, 0.05 mg/mL DNAse and 0.2 mg/mL lysozyme. The mixture was stirred at 4°C for 30 min then sonicated on ice. Lysate was clarified by centrifugation (F14-14×50cy rotor, 12,000 rpm, 20 min, 4°C) and then filtering through a 0.45 μm syringe filter. Affinity purification was done using a running buffer with 20 mM Tris (pH 7.5) and 150 mM KCl. TALON cobalt resin (Takara) was washed with water then equilibrated with 10 column volumes of running buffer. Clarified lysate was loaded on the column, washed with 10 column volumes of running buffer supplemented with 5 mM imidazole, then protein was eluted with running buffer supplemented with 50 mM imidazole. The nanobody was further purified using gel filtration with buffer containing 20 mM Tris (pH 7.5), 150 mM KCl, 0.5 mM EDTA. Peak fractions were then pooled and concentrated.

### MNT-002 Fab expression and purification

The MNT-002 antibody was generated by inserting the gene for the ShK peptide into the a β-ribbon ‘stalk’ of the ultralong CDR3 scaffold of a humanized bovine IgG as described in patent US10640574^62^. The Fab version of the antibody fusion (Fab-ShK) was expressed in freestyle HEK293 (293F) cells (Thermo Scientific) following the manufacturer’s protocol. In brief, 293F cells were seeded at a density of 1.0×10^6^ per ml prior to transfection. Cells were transfected with 293fectin (Thermo Scientific) combined with pFUSE-based plasmids (InvivoGen) encoding both heavy (HC) and light chains (LC) (HC:LC ratio of 1:1). Cell culture supernatant containing expressed protein was collected five days post-transfection, filtered through a 0.45 μm PES filter and concentrated at 4°C using an Amicon Ultra-15 centrifugal filter unit (molecular weight cut-off of 10,000 Da) (Millipore Sigma). Protein was further purified using CaptureSelect CH1-XL affinity resin (Thermo Scientific) following the manufacturer’s protocol. Purified protein was eluted in 50 mM sodium acetate (pH 4.0), then concentrated and buffer-exchanged into 20 mM Tris (pH 7.5) with 150 mM KCl and 0.5 mM EDTA at 4°C using Amicon Ultra-4 centrifugal filter unit (molecular weight cut-off of 10,000 Da) (Millipore Sigma). The Fab yield was quantified by measuring absorbance at 280 nm on a Nanodrop One (Thermo Scientific) and evaluated by SDS-PAGE under reducing and non-reducing conditions.

### Electrophysiology

The cDNA encoding the Kv1.3 expression construct used for protein production and cryo-EM was cloned into a pGEM vector^63^, linearized with NheI and transcribed by using a mMESSAGE mMACHINE™ T7 Transcription Kit (Invitrogen/ThermoFisher). Female *Xenopus laevis* animals were housed, and surgery was performed according to the guidelines of the National Institutes of Health, Office of Animal Care and Use (OACU) (Protocol Number 1253–18). Oocytes were removed surgically and incubated with agitation for 1 hr in a solution containing (in mM) 82.5 NaCl, 2.5 KCl, 1 MgCl_2_, 5 HEPES, pH 7.6 (with NaOH), and collagenase (1.2 mg/ml; Worthington Biochemical, Lakewood, NJ). Defolliculated oocytes were injected with 50 nl of channel RNA (~500 ng/ml) and maintained at 17°C for 1-3 days in an ND96 oocyte maintenance buffer, containing (in mM): 96 NaCl, 2 KCl, 5 HEPES, 1 MgCl_2_ and 1.8 CaCl_2_ plus 50 mg/ml gentamycin, pH 7.6 with NaOH. Voltage-clamp recordings were performed using the two-electrode voltage-clamp recording techniques (OC-725C; Warner Instruments) with a 150-μl recording chamber that was perfused continuously. Data were filtered at 1-3kHz and digitized at 10kHz using a Digidata 1321A interface board and pCLAMP 10 software (Axon; Molecular Devices). Microelectrode resistances were 0.2–0.8MΩ when filled with 3 M KCl. For recording macroscopic Kv channel currents, the external recording solution contained (in mM): 2 KCl, 98 NaCl, 5 HEPES, 1 MgCl_2_, and 0.3 CaCl_2_, pH 7.6, with NaOH. Fab-ShK or the nanobody were added to the control recording solution to the indicated concentrations. All experiments were performed at room temperature (22°C). Leak and background conductances were subtracted by arithmetically deducting the end of the tail pulse of each analyzed trace. Kv channel currents shown are non-subtracted.

G-V relationships were obtained by measuring tail currents at −50mV and a single Boltzmann function was fit to the data according to:

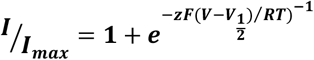

where z is the equivalent charge, V_1/2_ is the half-activation voltage, F is Faraday’s constant, R is the gas constant and T is temperature in Kelvin.

Time constants (τ) for nanobody binding and inactivation were obtained by fitting a single exponential function to the traces for depolarization between +20mV to +50mV. The traces were fitted from the beginning of the decaying phase to the end of the depolarizing trace using the following equation:

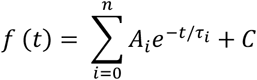

Sigmoidicity values for recovery from nanobody inhibition were obtained by fitting the following equation^64^ to the data obtained from the time-course experiments (**Fig. S5D**):

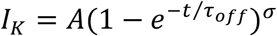

The amplitude of the sigmoidal curve A develops with a time constant Toff (ms), and sigmoidicity σ, which is unitless. When σ = 1, the equation describes a monoexponential rise, as would be expected from a process involving one transition. As σ increases the value of σ would indicate the number of transitions required to produce such a sigmoidicity.

### Cryo-EM sample preparation and data acquisition

Samples were prepared using human Kv1.3 protein (3 mg/mL) without a binding partner, or with nanobody at 1.3 mg/mL (3:1 molar ratio, nanobody:Kv1.3 subunit), or with Fab-ShK at 0.4 mg/mL (1:1 molar ratio, Fab-ShK:Kv1.3 tetramer). UltrAuFoil 1.2/1.3 300 mesh grids (Quantifoil) were plasma treated and vitrified samples were prepared by adding a 2.5 μL droplet of sample solution to a grid, then blotting (2 sec blot time, 0 to −4 blot force range) and plunge-freezing in liquid ethane using a Vitrobot Mk IV (Thermo Fisher).

Single particle images were collected with a Titan Krios electron microscope (Thermo Fisher) operated at 300 kV and a nominal magnification of 105,000 × and equipped with a K3 camera (Gatan) set in superresolution mode (0.4260 Å pixel size). Leginon was used for automated collection of images with ice thickness ranging between 20 nm and 120 nm^65,66^. Movies were collected at nominal defocus values of ~1.0-2.0 μm and dose-fractionation into 48 frames with a total exposure time of 2.4 sec and total dose of ~51 e-Å^-2^.

### Image processing and structure analysis

Movie stacks were corrected for beam-induced motion with two-fold binning and dose-weighted in Relion^48^. The resulting images were used for contrast transfer function (CTF) estimation with CTFFIND4.1^67^. Datasets were then processed as follows.

For the unbound Kv1.3 dataset an initial particle set was picked with the reference-free Laplacian-of-Gaussian (LoG) tool in Relion and particles were extracted with a box size of 320 and downscaled to 128. Particles were imported into cryoSPARC and processed with 2D classification, ab initio reconstruction and heterogeneous refinement to generate a well-defined particle subset. The subset was re-extracted in Relion using a box size of 320 and the 3D model generation tool in Relion was used to create a template map for particle picking. Particle picking was done with an interparticle distance of 150 Å to avoid picking duplicates and yielded 526,276 particles. The new particle set was extracted with a box size of 320 and imported into cryoSPARC. Particles were processed with ab initio reconstruction and multiple rounds of heterogeneous refinement to isolate a subset of 190,793 particles which were refined with homogeneous and non-uniform refinement to 3.30 Å with C1 symmetry. Particle polishing was done in Relion and ab initio reconstruction, homogeneous refinement and non-uniform refinement were done in cryoSPARC to obtain a 3.12 Å map with C1 symmetry. A final non-uniform refinement with C4 symmetry was used to generate a map with 2.89 Å resolution.

The Kv1.3 dataset was further processed to isolate maps for the D1 and D2 subunit conformations. The particle set used to generate the C4 symmetric tetramer map (190,793 tetramer particles) was subjected to symmetry expansion in Relion to generate images of individual tetramer subunits (763,172 subunit particles). The subunit images were processed with multiple rounds of 3D classification without alignment in Relion using a mask around a single subunit. Uninterpretable classes or classes with a mixture of D1/D2 loop conformations were discarded, and D1 and D2 subunit maps were resolved with 107,045 and 217,344 particles, respectively. These particles were used for additional analysis on tetramer compositions as presented in Table S2.

For the Kv1.3 with nanobody dataset an initial particle set was picked with the LoG tool in Relion and particles were extracted with a box size of 416 and downscaled to 128. Particles were imported into cryoSPARC and approximately half of the LoG-picked particles were randomly selected and processed with 2D classification, ab initio reconstruction and heterogeneous refinement to generate a well-defined subset of 27,427 particles. The subset was re-extracted in Relion using a box size of 416 and the 3D model generation tool in Relion was used to create a template map for particle picking. Particle picking was done with an interparticle distance of 150 Å to avoid picking duplicates and yielded 394,303 particles. The new particle set was extracted with a box size of 416 and imported into cryoSPARC. Particles were processed with multiple rounds of heterogeneous refinement to isolate a subset of 138,728 particles which were refined with homogeneous and non-uniform refinement to 3.62 Å with C4 symmetry. Particle polishing was done in Relion and ab initio reconstruction and heterogeneous refinement were done in cryoSPARC to isolate a subset of 123,722 polished particles. These particles were further refined with homogeneous refinement and non-uniform refinement to generate a final map at 3.25 Å resolution map with C4 symmetry.

For the Kv1.3 with Fab-ShK dataset an initial particle set was picked with the LoG tool in Relion, extracted with a box size of 440 and downscaled to 128, and processed with 2D classification in cryoSPARC to identify an interpretable subset. Templates were generated with 2D classification in Relion and used for particle picking to yield 482,405 particles. The particles were extracted with a box size of 440, imported into cryoSPARC, and 2D classification was used to isolate a subset of 447,444. Particles were subjected to iterative heterogeneous refinement in cryoSPARC to isolate 90,267 particles that were used for nonuniform refinement to generate a 3.71 Å map. Kv1.3 and the ShK toxin were well-defined in the map, but only the overall molecule profile was visible for the Fab region. Attempts to improve the Fab region with local refinement were unsuccessful. The particles were subjected to particle polishing in Relion and then refined in cryoSPARC with non-uniform refinement to a final resolution of 3.39 Å. C1 symmetry was used throughout.

### Structural modeling

A homology model for Kv1.3 was generated using PDB 2R9R (Kv1.2 crystal structure). The homology model was docked into each Kv1.3 map using Chimera^68^ and the model for each structure refined using Coot^69^. For the unbound Kv1.3 structure, separate models were built for the D1 and D2 conformations. This was done by using the D1 or D2 subunit maps in parallel with the tetrameric C4 symmetric consensus map. The two models are virtually identical except for the selectivity filter region. For the Kv1.3 with nanobody A0194009G09 structure a homology model was generated using PDB 6V80, docked into the map using Chimera, and refined using Coot. Separate versions of the Kv1.3-NB complex were modeled for the D1 and D3 conformations, which varied differed only in the rotamer orientation of Tyr447 and associated local changes in the protein backbone. For the map of Kv1.3 with the Fab-ShK fragment from MNT-002, a homology model was generated from PDB 4LFQ (ShK) and PDB 6OO0 (bovine Fab) and docked into the map using Chimera. The ShK toxin model was refined using Coot. The Fab region of the map was sufficient to orient the Fab but was not sufficient for further modeling so all residues were stubbed to alanine. The extracellular VSD linkers were not resolved so were not modeled. The exception to this is the first segment of the S1-S2 loop which was stabilized by nanobody binding in the Kv1.3-nanobody complex so could be modeled.

## DATA AVAILABILITY

The cryo-EM density maps and models for Kv1.3, Kv1.3 with nanobody A0194009G09, and Kv1.3 with the Fab-ShK from MNT-002 have been deposited in the Protein Data Bank (PDB) and Electron Microscopy Data Bank and will be released upon publication. The PDB accession codes are 7SSX (Kv1.3 D1), 7SSY (Kv1.3 D2), 7SSZ (Kv1.3 with nanobody D1), 7ST0 (Kv1.3 with nanobody D3) and 7SSV (Kv1.3 with Fab-ShK) and the EMD accession codes are EMD-25416 (Kv1.3 map, D1 subunit map, D2 subunit map), EMD-25417 (Kv1.3 with nanobody) and EMD-25414 (Kv1.3 with Fab-ShK).

