## supplemental_figures for "Structures of the T cell potassium channel Kv1.3 with immunoglobulin modulators"

### SUPPLEMENTAL FIGURES AND TABLES

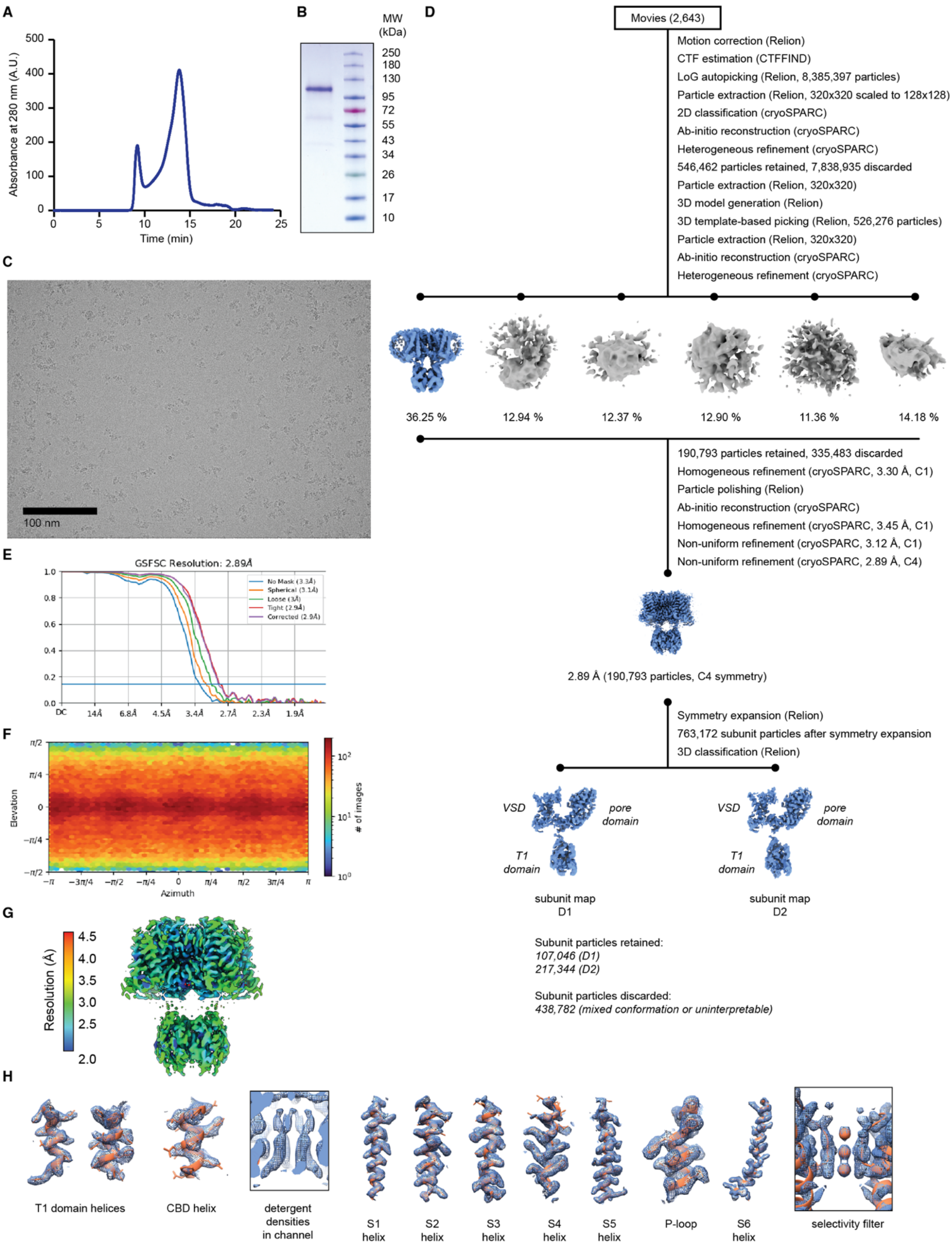

**Supplemental Figure 1. Purification, cryo-EM, and structure determination for human Kv1.3.** **(A-B)** Gel filtration trace (A) and SDS-PAGE (B) for purified human Kv1.3. **(C)** Representative cryo-EM image of Kv1.3. **(D)** Data processing workflow for Kv1.3. Cryo-EM density maps are color-coded as blue or gray according to whether they are retained or discarded, respectively. The percentage of particles in each class from heterogeneous refinement is given. The final two maps are for individual Kv1.3 subunits in the D1 (left) and D2 (right) conformations isolated by symmetry expansion and 3D classification. **(E)** Fourier shell correlation (FSC) curves as generated by cryoSPARC. **(F)** Angular distribution plot for particles in the reconstruction as generated by cryoSPARC. **(G)** Local resolution heat map. Data was processed with C1 and C4 symmetry as indicated, with the final map refined with C4 symmetry. **(H)** Select regions of the cryo-EM density map highlighting the T1 domain, CBD helix, presumed detergent densities observed in the channel, P-loop, S1-S6 helices, and the selectivity filter.

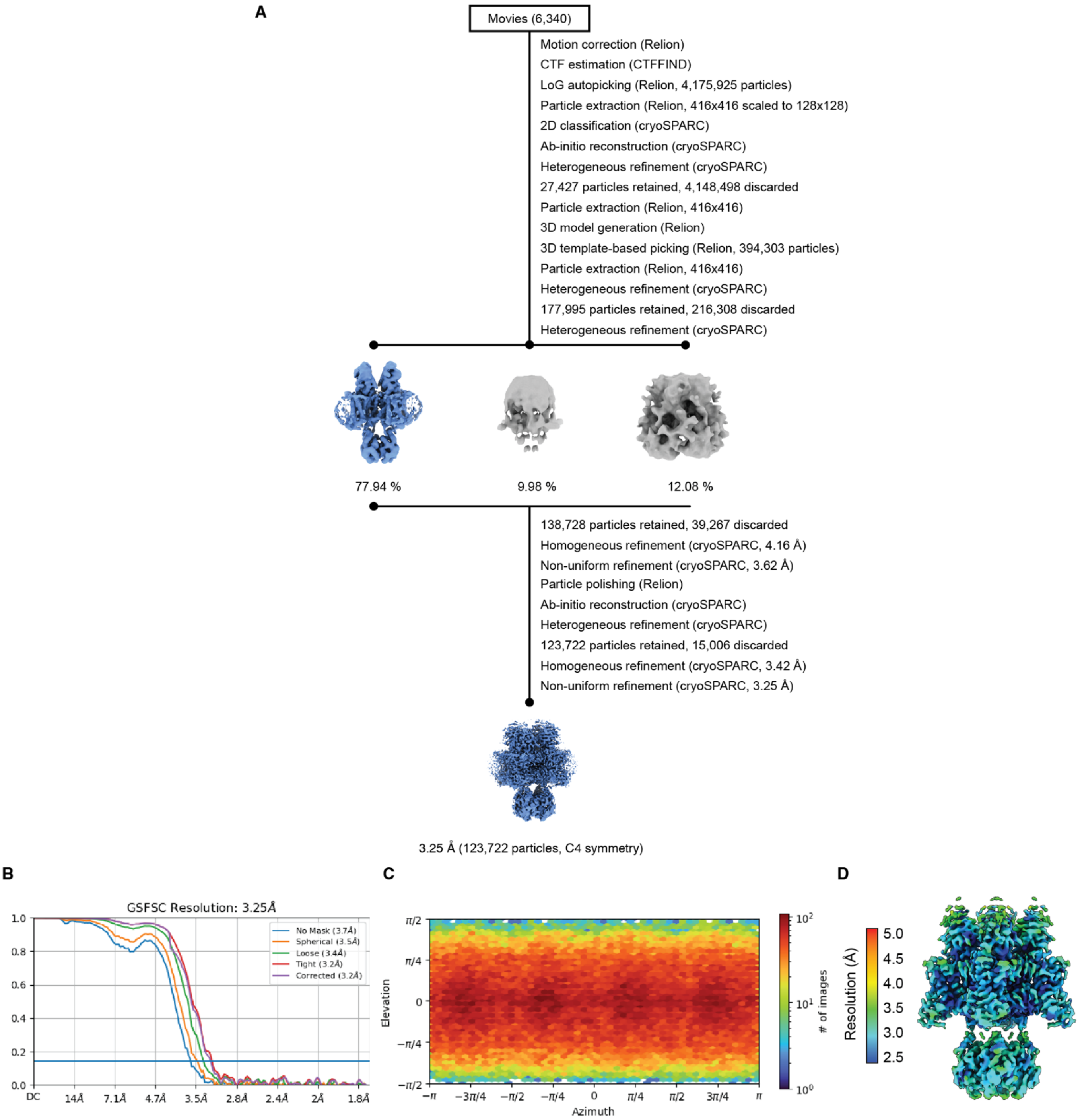
A
Movies (6,340)
Motion correction (Relion)
CTF estimation (CTFFIND)
LoG autopicking (Relion, 4,175,925 particles)
Particle extraction (Relion, 416x416 scaled to 128x128)
2D classification (cryoSPARC)
Ab-initio reconstruction (cryoSPARC)
Heterogeneous refinement (cryoSPARC)
27,427 particles retained, 4,148,498 discarded
Particle extraction (Relion, 416x416)
3D model generation (Relion)
3D template-based picking (Relion, 394,303 particles)
Particle extraction (Relion, 416x416)
Heterogeneous refinement (cryoSPARC)
177,995 particles retained, 216,308 discarded
Heterogeneous refinement (cryoSPARC)
77.94 %
9.98 %
12.08 %
138,728 particles retained, 39,267 discarded
Homogeneous refinement (cryoSPARC, 4.16 Å)
Non-uniform refinement (cryoSPARC, 3.62 Å)
Particle polishing (Relion)
Ab-initio reconstruction (cryoSPARC)
Heterogeneous refinement (cryoSPARC)
123,722 particles retained, 15,006 discarded
Homogeneous refinement (cryoSPARC, 3.42 Å)
Non-uniform refinement (cryoSPARC, 3.25 Å)
3.25 Å (123,722 particles, C4 symmetry)
B
GSFSC Resolution: 3.25Å
No Mask (3.7Å)
Spherical (3.5Å)
Loose (3.4Å)
Tight (3.2Å)
Corrected (3.2Å)
1.0
0.8
0.6
0.4
0.2
0.0
DC
14Å
7.1Å
4.7Å
3.5Å
2.8Å
2.4Å
2Å
1.8Å
C
Elevation
π/2
π/4
0
−π/4
−π/2
Azimuth
−π
−3π/4
−π/2
−π/4
0
π/4
π/2
3π/4
π
### of images
$10^2$
$10^1$
$10^0$
D
Resolution (Å)
5.0
4.5
4.0
3.5
3.0
2.5

**Supplemental Figure 2. Structure determination for human Kv1.3 with nanobody A0194009G09.** **(A)** Data processing workflow for Kv1.3 in complex with A0194009G09 nanobodies. Cryo-EM density maps are color-coded as blue or gray according to whether they are retained or discarded, respectively. The percentage of particles in each class from heterogeneous refinement is given. **(B)** Fourier shell correlation (FSC) curves as generated by cryoSPARC. **(C)** Angular distribution plot for particles in the reconstruction as generated by cryoSPARC. **(D)** Local resolution heat map. Data was processed with C4 symmetry.

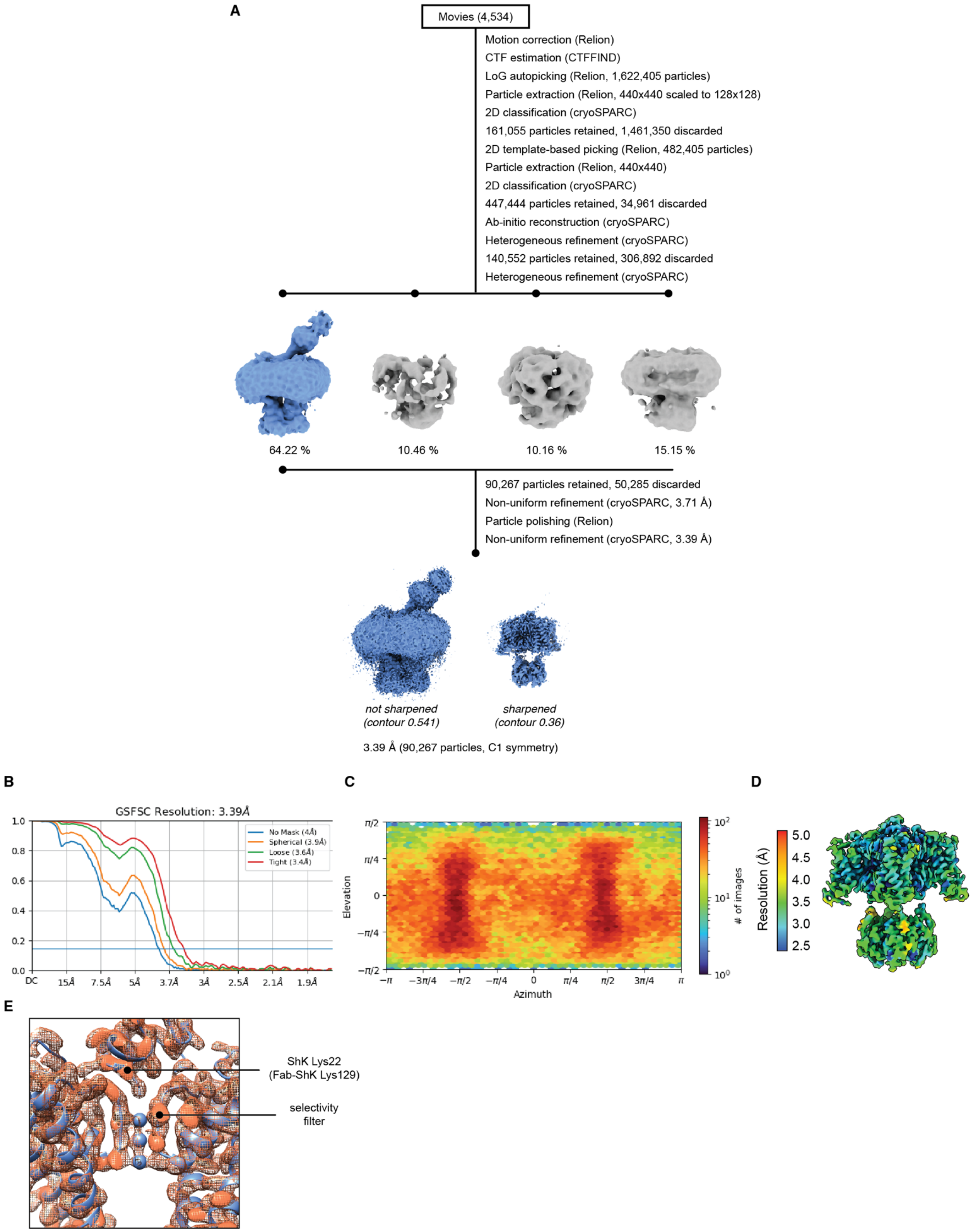
A
Movies (4,534)
Motion correction (Relion)
CTF estimation (CTFFIND)
LoG autopicking (Relion, 1,622,405 particles)
Particle extraction (Relion, 440x440 scaled to 128x128)
2D classification (cryoSPARC)
161,055 particles retained, 1,461,350 discarded
2D template-based picking (Relion, 482,405 particles)
Particle extraction (Relion, 440x440)
2D classification (cryoSPARC)
447,444 particles retained, 34,961 discarded
Ab-initio reconstruction (cryoSPARC)
Heterogeneous refinement (cryoSPARC)
140,552 particles retained, 306,892 discarded
Heterogeneous refinement (cryoSPARC)
64.22 %
10.46 %
10.16 %
15.15 %
90,267 particles retained, 50,285 discarded
Non-uniform refinement (cryoSPARC, 3.71 Å)
Particle polishing (Relion)
Non-uniform refinement (cryoSPARC, 3.39 Å)
not sharpened (contour 0.541)
sharpened (contour 0.36)
3.39 Å (90,267 particles, C1 symmetry)
B
GSFSC Resolution: 3.39Å
No Mask (4Å)
Spherical (3.9Å)
Loose (3.6Å)
Tight (3.4Å)
DC
15Å
7.5Å
5Å
3.7Å
3Å
2.5Å
2.1Å
1.9Å
C
Elevation
Azimuth
### of images
D
Resolution (Å)
5.0
4.5
4.0
3.5
3.0
2.5
E
ShK Lys22 (Fab-ShK Lys129)
selectivity filter

**Supplemental Figure 3. Structure determination for human Kv1.3 with Fab-ShK from the MNT-002 antibody.**

**(A)** Data processing workflow for Kv1.3 in complex with Fab-ShK from the MNT-002 antibody. Cryo-EM density maps are color-coded as blue or gray according to whether they are retained or discarded, respectively. The percentage of particles in each class from heterogeneous refinement is given. The final map is shown without sharpening and at low contour to visualize the Fab density, and with sharpening and at high contour to visualize high-resolution structural features. **(B)** Fourier shell correlation (FSC) curves as generated by cryoSPARC. **(C)** Angular distribution plot for particles in the reconstruction as generated by cryoSPARC. **(D)** Local resolution heat map. Data was processed with C1 symmetry. **(E)** Kv1.3 map with model for the selectivity filter and ShK region of the structure. The structure highlights the potassium ions and key lysine residue of ShK that is coordinated by the selectivity filter. The lysine is position 22 in ShK or position 129 in the Fab-ShK fusion. The front and rear Kv1.3 subunits are omitted for clarity.

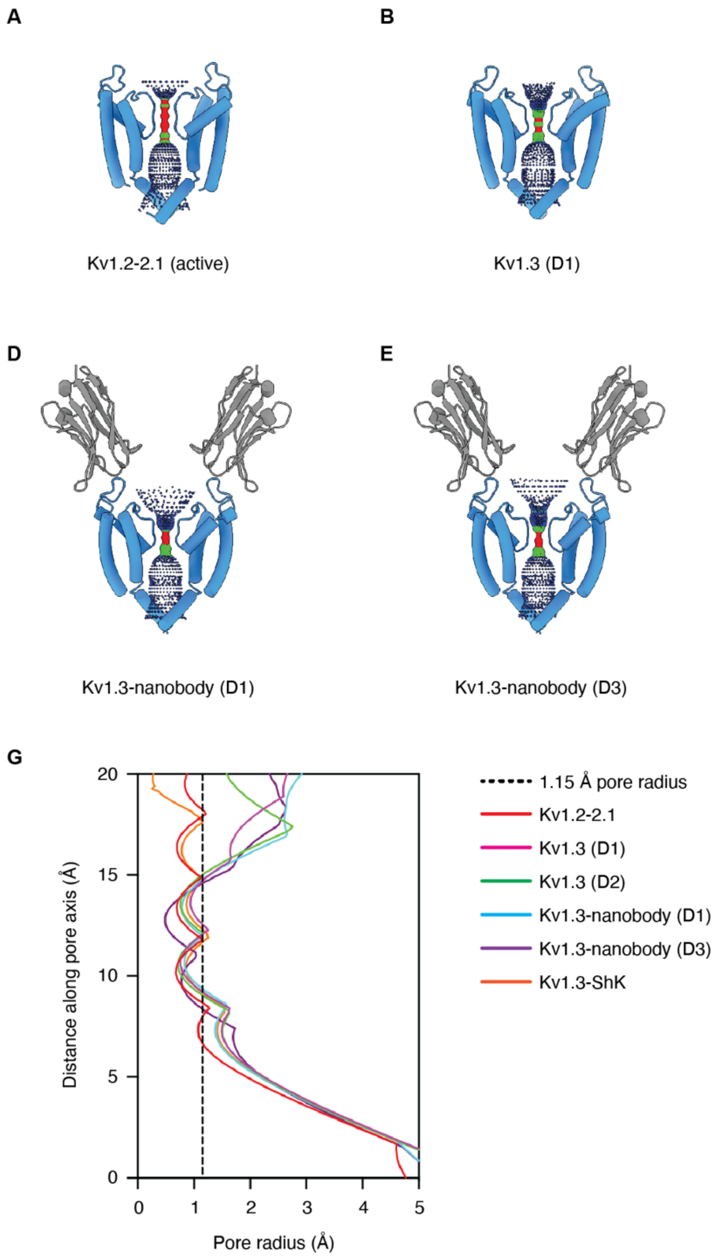

**Supplemental Figure 4. Visualization of pore profiles.**

**(A-F)** Visualizations of the pore structures for Kv1.2-2.1 (A), Kv1.3 in D1 conformation (B), Kv1.3 in D2 conformation (C), Kv1.3 with nanobodies in D1 conformation (D), Kv1.3 with nanobodies in D3 conformation (E), and Kv1.3 with ShK (F). Each structure is shown with only two subunits for visual clarity. Panels (D) and (E) show the Kv1.3 pore with two nanobodies, and panel (F) shows the Kv1.3 pore with the ShK toxin. **(G)** Plot of pore radius along the length of each pore structure in (A-F).

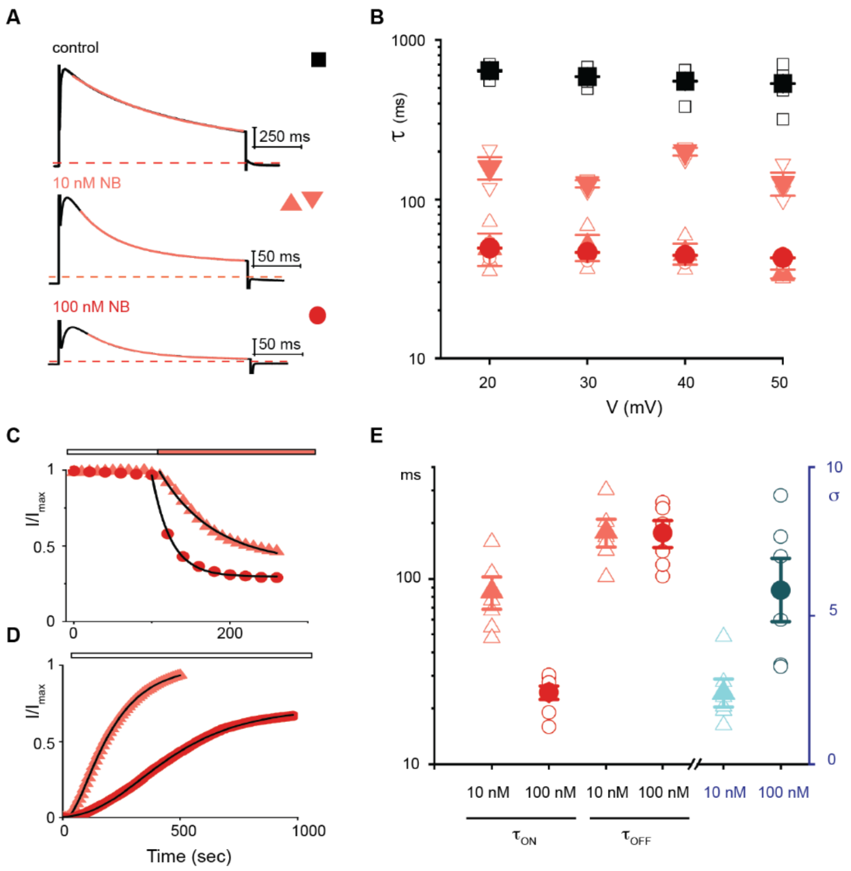

**Supplemental Figure 5. Concentration-dependence for acceleration of inactivation by the NB.**
**(A)** Representative Kv1.3 current traces obtained by depolarization to +50 mV when eliciting a family of traces by stepping from -80 mV to +50 mV in control conditions (top panel), in the presence of 10 nM NB (middle panel) and 100 nM NB (bottom panel). Holding voltage was -80 mV and tail voltage was -50 mV. Red dotted lined denotes zero current level. The control trace and that for 100 nM NB are from one cell and the trace for 10 nM NB is from another cell. Solid orange lines are fits of single (control and 100 nM) or double exponential (10 nM) functions to the current traces (see Methods). Current scale bars are 1 μA. **(B)** Influence of the NB on inactivation for varying test depolarizations. For control, mean values are shown as filled black squares and individual measurements as open black squares (n=6). For 10 nM NB, mean values for $\tau_1$ are shown as filled upward facing orange triangles and those for $\tau_2$ filled as downward facing orange triangles, with individual measurements as open triangles (n=3). For 100 nM NB mean values are shown as filled orange circles, with individual measurements as open circles (n=3). In all instances, error bars are S.E.M. **(C)** Time courses for onset of inhibition by the NB. Peak currents were measured during 500 ms pulses to +40 mV from a holding potential of -80 mV and normalized to the maximum value before the addition of NB (orange box) at a concentration of 10 nM (orange triangles)

and 100 nM (orange circles). Time scales were synchronized for plotting purposes. Solid black lines are fits to a single exponential function to the time course (see Methods). **(D)** Time course for recovery from inhibition by the NB. Peak currents were measured during 500 ms pulse to +40 mV from a holding potential of -80 mV following removal of the NB (white box). Currents were scaled by setting the initial values in NB to zero and normalizing to the maximum value before the addition of the NB and to facilitate visualization and fitting. 10 nM NB (filled orange triangles) and 100 nM (filled dark orange circles). Solid black lines are fits to a sigmoidal function to the individual time courses (see Methods). **(E)** Concentration dependence for onset of ($\tau_{ON}$) and recovery from ($\tau_{OFF}$) inhibition by the NB. For 10 nM NB, mean values for $\tau_{ON}$ and $\tau_{OFF}$ are shown as filled orange triangles, and $\sigma$ as light blue triangles with individual measurements as open triangles. For 100 nM NB, mean $\tau_{ON}$ and $\tau_{OFF}$ values are shown as filled dark orange circles and $\sigma$ as dark blue circles, with individual measurements as open circles. $\tau_{ON}$ values were obtained from the single exponential fits to the time courses as shown in panel C. $\tau_{OFF}$ and $\sigma$ values were obtained from the sigmoidal fits to the time courses as shown in panel (D). In all instances, n = 6 and error bars are S.E.M.

| | Kv1.3 apo | Kv1.3 with A0194009G09 nanobody | Kv1.3 with MNT-002 Fab |
|---|---|---|---|
| **Microscope** | Krios | Krios | Krios |
| **Magnification** | 105,000 | 105,000 | 105,000 |
| **Voltage (kV)** | 300 | 300 | 300 |
| **Electron exposure (e–/Å^2)** | 53.7 and 54.23 | 50.93 and 55.97 | 51.36 |
| **Defocus (μm)** | 1.3-2.0 | 1.3-2.0 | 1.3-2.0 |
| **Pixel size (Å)** | 0.426 | 0.426 | 0.426 |
| **Binned pixel size (Å)** | 0.852 | 0.852 | 0.852 |
| **Frames/movie (#)** | 48 | 48 | 48 |
| **Symmetry imposed** | C4 | C4 | C1 |
| **Movies (#)** | 2,643 | 6,340 | 4,534 |
| **Particle images (#)** | 190,793 | 123,722 | 90,267 |
| **Box size (pixels)** | 320 | 416 | 440 |
| **Map resolution (Å)** | 2.89 | 3.25 | 3.39 |
| **FSC threshold** | 0.143 | 0.143 | 0.143 |

**Supplemental Table 1. Cryo-EM data collection and image processing.**

Information on cryo-EM data collection and image processing for Kv1.3 structures reported in the study.

| | |
|---|---|
| **Tetramers used for symmetry expansion** | 190,793 |
| **Subunits generated by symmetry expansion** | 763,172 |
| **Subunits classified as D1** | 107,046 |
| **Subunits classified as D2** | 217,344 |
| **Subunits in conformationally mixed or uninterpretable classes (discarded)** | 438,782 |
| **Tetramers with all 4 subunits identified** | 6,598 |
| **Tetramers with 1,2 or 3 subunits not identified (discarded)** | 184,195 |
| **D1:D2 ratios in tetramers with all 4 subunits identified:** | |
| *0:4* | 1,752 tetramers (26.55%) |
| *1:3* | 2,032 tetramers (30.8%) |
| *2:2* | 1,618 tetramers (24.52%) |
| *3:1* | 900 tetramers (13.64%) |
| *4:0* | 296 tetramers (4.49%) |
| **D1 subunits analyzed** | 9,152 (34.68%) |
| **D2 subunits analyzed** | 17,240 (65.32%) |

**Supplemental Table 2. Conformational analysis of unbound Kv1.3.**

Statistics from conformational analysis of unbound Kv1.3. The set of tetramer particles used as input for the analysis is the same as used for in the reported tetramer structure and contains 190,793 particles (**Fig. S1**).
